# Dynamics of single cell-cell junctions as an indicator of cell state switch

**DOI:** 10.64898/2026.02.18.706725

**Authors:** Kavya Senthilazhagan, Amit Das

## Abstract

Cell-cell junctions (CCJs) are dynamic biopolymeric systems essential for adherence of biological cells organizing into a tissue or organ. In vertebrate organisms, CCJs present in stable epithelial tissues are maintained primarily by cell-to-cell protein bridges made of an adhesion receptor E-cadherin. CCJs are destabilized during Epithelial (E) to mesenchymal (M) transformation (EMT), an essential step in cancer metastasis, where cells switch states and acquire migratory features. An essential trigger for EMT is a cadherin switch process from E-cadherins with N-cadherin, another cadherin isoform. EMT proceeds through several intermediate states referred to as hybrid E/M cells. These states are characterized by mixed levels of E- and N-cadherins at the junctions and exhibit versatile cancerous traits that are more aggressive than cancerous fully mesenchymal cells. As a result, many such states have emerged as key targets in cancer therapy. However, development of a therapeutic design to counter the hybrid E/M cells has been limited by the absence of a comprehensive understanding of the mechanics and dynamics of hybrid E/M states. Here, we develop a physical model of CCJs as a non-equilibrium system composed of a variable ratio of E- and N-cadherins, considered as coarse-grained molecules driven by ATP-powered machinery observed at CCJs. Our model predicts a robust measure of strength of junctions that captures previous experimental observations, and reveals a minimal mechanochemical landscape of hybrid E/M states. We show the emergence of several groups of CCJ states in this landscape with variable adhesion strengths, many of which resemble different hybrid E/M-characteristics observed experimentally. Finally, we identify that a difference in mechanosenstivity of the two cadherin isoforms towards cytoskeletal forces could be why the hybrid E/M states come into existence.

## I. INTRODUCTION

Adherence of similar cells is essential for development of multicellular organisms. The interfaces of two cells in proximity adhere to form a region known broadly as the ‘cell-cell junctions’ (CCJs) which allow the cells organize into cohesive, functional tissues [1]. The CCJs are biopolymeric assemblies that exhibit strong non-equilibrium nature as they can remain stable for long periods of time (hours to days or even years) [2] or undergo dynamic remodeling under specific conditions [3]. The diverse mechanical states of CCJ’s have been compared with both fluidized or glassy systems during morphogenesis of vertebrate organisms [4].

The core mechanism for formation of CCJs is orchestrated by *trans* linkages formed by a variety of adhesion receptors, which are cell-to-cell protein-bridges [1]. These linkages are primarily homotypic in nature, comprising of many molecules of E-cadherin, a highly abundant member of the cadherin receptor family in vertebrates. However, CCJs are well-known to conditionally exhibit mixed-populations of cadherins to facilitate transformation between stable tissue states or ‘phenotypes’. One such important state-switch is EMT [5, 6], a necessary mechanism during embryonic morphogenesis, wound-repair and cancer progression.

A systematic decrease of E-cadherin levels has been recently pinpointed [7] as an initiation signal for EMT, leading to a loss of their *trans* clustering. This is accompanied by the upregulation of a different cadherin isoform, namely N-cadherin, governed by specific transcription factors and epigenetic pathways [8]. N-cadherin, albeit having a similar sequence and structure to E-cadherin, offers different and weaker homotypic linkages compared to E-cadherin [9–12]. The down- and upregulation of E- and N-cadherin, respectively, however, happen smoothly that renders EMT a continuous transformation between the E- and the M-states. Thus, EMT is not a binary switch. The landscape of EMT has been proposed to be decorated with several metastable and stable, intermediate ‘hybrid E/M’ states [5, 6, 13].

The hybrid E/M phenotype is implicated to be the ‘fittest’ for metastasis among different cancerous phenotypes and main culprits for the growth of secondary tumors [14]. They are more tumorigenic compared to fully M cells and resistant to therapy [15–17]. Thus, potential therapeutic targets may exist in the EMT spectrum, in the form of different hybrid E/M cell populations [18, 19]. However, we don’t fully understand the EMT spectrum in a way that allows us to identify specific hybrid E/M populations through a measurable cellular metric. This type of identification may be the first step to developing a therapeutic strategy to counter these metastatically potent groups of cells. From a physics point of view, these junctions could represent a new type of adaptable, non-equilibrium system with unique mechanical signatures, capable of diverging towards any terminal state on the EMT spectrum.

Most of the existing approaches to elucidate the EMT landscape has been focused on the biochemical signaling networks governing EMT [16]. A large body of mathematical modeling work exists from Jolly, Levine and co-workers in this direction based on gene regulatory circuits [20–23]. Conversely, recent experimental studies have used morphological traits, like changes in cell shape, nuclear shape, cytoskeletal architecture to understand the progression of EMT [24– 28]. Our recent work on drug-induced EMT in lung epithelial cells [25] revealed clear evidence that hybrid E/M cells manifest a strongly heterogeneous distribution of E- and N-cadherin on the junctions and a more flexible migratory phe-notype compared to mature epithelial tissues. The local patterning of the cadherin clusters and consequent mechanical heterogeneity of CCJs could be a controlling agency behind such flexibility. However, a comprehensive understanding of the connection between the biochemical signatures of EMT and local mechanics of CCJs at the molecular scale is still completely lacking.

Existing physical models on CCJs have concentrated either on a pure mechanical picture of the entire junction [4] or a detailed molecular description at a very limited scale in comparison to their spatial scale [12, 29]. However, all these studies lack a vital physiological source of non-equilibrium character of E-cadherin dynamics in CCJs - recycling of the protein molecules that limits the accumulation of extremely big cadherin clusters [30]. In the current work, we build a physical model of CCJs considering the cadherins as coarse-grained spherical molecules, thus preserving the molecular-scale features. The cadherins in the model explore a spatial scale of several microns comparable to that of the length of a single junction. This way our model captures the overall features of the junctions as well. We consider various essential ingredients of cadherin dynamics, including the active cortical forces on the cadherins as well as recycling between membrane and cytoplasmic populations.

We quantify the strength of individual CCJs as the main EMT indicator. We consider two essential biological controllers of the junction strength associated with EMT. First one is the biochemical controller: the ratio of N-to E-cadherin densities on the cell surface. This is because the hybrid E/M cells are characterized with a variable E- and N-cadherin expression patterns and exhibit weaker junctions compared to fully epithelial cells [5, 8, 31]. The second factor is the mechanial controller: the cortical active forces from the actin cytoskeleton driven by myosin motors. E-cadherin *trans*-linkages are mechanically stabilized via a catch bond effect orchestrated by cadherin-actin interactions [32]. Lifetimes of such linkages increase as they are pulled by mechanical forces up to a threshold. Furthermore, the lateral dynamics of cortical actin filaments promotes formation of cadherin *cis*-clusters by facilitating nucleation events [33]. These *cis* clusters also stabilize junctions through a cooperative engagement with the *trans* linkages [9, 34]. Recent evidence hint that E- and N-cadherin possibly experience such cortical forces differently that leads to their dissimilar adhesive strength [11, 12]. We incorporate these effects in form of a species-specific, lateral active propulsive force that is added on the diffusing cadherin molecules. The exact form of this active force is inspired by extremely well studied active Brownian particles [35]. We consider a stronger *trans*-interaction strength that *cis* to capture the essence of catch bond effect, which breaks down only when activity is significantly high.

We validate the physical model by capturing the steadystate clustering behavior reported previously for junctional E-cadherin molecules in Drosophila embryos [33]. Our model recapitulates the growth dynamics of junctions and the statistical distribution of cluster sizes. Subsequently, we identify an appropriate measure of junction strength in terms of the adhesion energy per unit interfacial area and recover a physiologically relevant range of values for this quantity. Finally, we predict a minimal landscape of different junctional states associated with EMT, where we link specific ones to different hybrid E/M states and propose experimental strategies for their identification. We predict that the above-mentioned difference in the adhesive strength of E- and N-cadherin due to their differential interaction to cortical forces constitutes a probable mechanism for emergence of the hybrid E/M states.

## II. DETAILS OF THE MODEL

### A. System

We coarse-grain all the cadherin extracellular domains as single spheres of diameter σ, a uniform average diameter for E- or N-cadherin (Fig. 1a). These molecules are constrained to diffuse on a square flat surface representing a cell membrane patch of lateral dimension *L*, determined by the surface density of cadherin molecules. We bring two such flat surfaces in close proximity so that molecules from both the membranes can form *trans* linkages (Fig. 1a, b). We apply periodic boundary conditions only along X and Y directions to simulate effects of a much bigger membrane. Thus, we explicitly consider the two layers of junctional cadherin molecules (Fig. 1b), unlike recent pseudo-bilayer approaches [36, 37] where projections of E-cadherin molecules and F-actin on a single plane were considered. Through the explicit approach, we capture the effects of physical crowding on the lateral movements of the molecules at the cell surface. The extracellular space, separating two surfaces making up a junction, typically spans 15-30 nm [38]. Here, we maintain a constant distance of σ(≈6 *nm*)between the two surfaces to ensure *trans*-contacts (Fig. 1a).

**FIG. 1.**
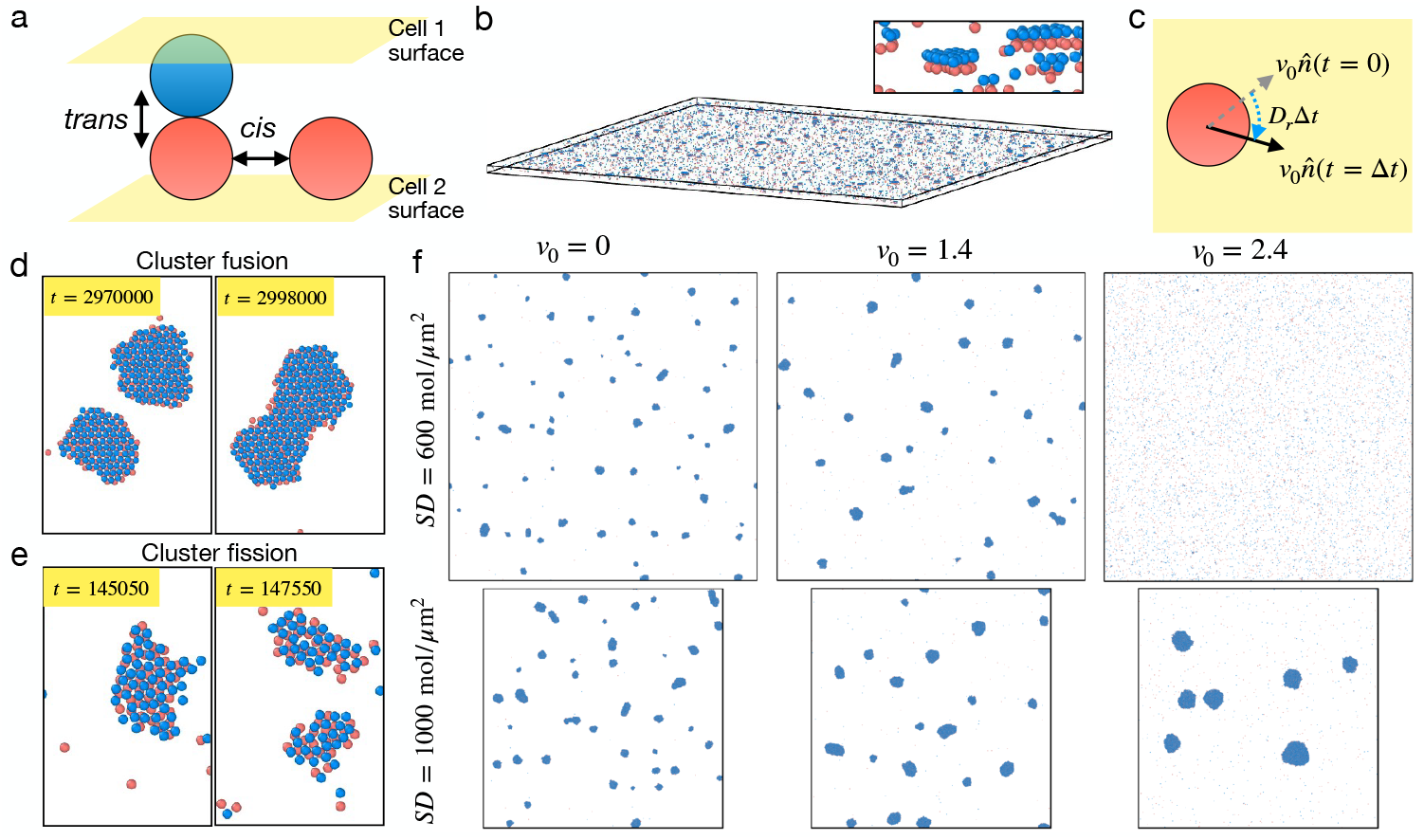
(a) Schematic of model cadherins and their *cis* and *trans* interactions. (b) A representative snapshot of the simulated system with a zoomed view of a few clusters shown in the inset. (c) Implementation of active propulsion on cadherins due to F-actin. Representative examples of (d) fusion of two clusters into one and (e) a single cluster into two. The system shown in (e) is at a higher activity, *v*_0_ = 2.4, compared to that in (d) with *v*_0_ = 1.4. (f) Effects of *v*_0_ and surface density (*SD*) on cluster formation. Top-row, at *SD* = 600 molecules*/µm*^2^, the cluster formation enhances initially with increasing *v*_0_ but then gets suppressed for higher *v*_0_. Bottom-row, at *SD* = 1000 molecules*/µm*^2^, the above trend is replaced by a monotonic increase in cluster formation as *v*_0_ increases. These indicate that activity can help or resist clustering only up to a certain surface density. Beyond a threshold *SD*, density-driven equilibrium clustering mechanisms takes over.

### B. Energetics

We describe all intermolecular interaction energies via a Lennard-Jones potential function of the form: 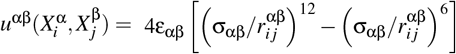 which describes the pair-wise interaction between the *i*th molecule of the species α with *j*th molecule of the species β (α, β = *E, N*). We consider, 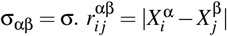 represents the scalar distance between molecules *i* of species α and *j* of species β.

We tune the strength of interaction ε_αβ_ between species α and β in units of thermal energy *k*_*B*_*T*, where *k*_*B*_ is the Boltzmann constant and *T* is the absolute temperature. Thus, ε_αβ_ *> k*_*B*_*T* would produce spontaneous attractive interactions among adjacent molecules and lead to clustering (Fig. 1d-f). We fix the values of ε_αβ_ based on literature. A significant number of previous studies have alluded to the *cis* and *trans*-dimerization of E-cadherin [9, 38–40]. Distinguishing between *cis* and *trans* complexes has been proven quite difficult because of their cooperative nature [34]. However, it has been consistently pointed out that strength of *trans* interaction is higher than that of *cis* [36, 40, 41]. Therefore, We choose a typical value for *trans*-ε_*EE*_ ≈ 4*k*_*B*_*T*, used previously [36], and keep a lower *cis*-ε_*EE*_ ≈ 2*k*_*B*_*T*. Recent micropipette aspiration [10], and AFM [12] studies *in vitro* have shown that N-cadherin has mechanically a much weaker *trans* interaction from E-cadherin. The comparison of E-E and N-N *cis*-interaction is, on the other hand, inconclusive with indirect suggestions that they are comparable. Accordingly, we set *cis*-ε_*NN*_ ≈ 2*k*_*B*_*T* and tune *trans*-ε_*NN*_ to study its effects on junction strength.

### C. Forces

Apart from intermolecular interactions, we consider three additional types of forces acting on each cadherin molecule: a frictional drag from the membrane, the stochastic Brownian forces from omnipresent random kicks from other membrane constituents, and the transient contractile stresses from cortical actin and myosin motors. The first is implicitly incorporated by a friction coefficient, Γ that we keep uniform for all molecules. The second is implemented by a Gaussian white noise with zero mean and variance set by the thermal energy and obeys the fluctuation-dissipation theorem. For the third, we focus only on the in-plane contribution of the actin-driven forces on the cadherins, as they constitute the predominant component as shown elsewhere [42–44]. We represented this contribution as a lateral propulsive force of fixed strength 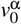, applicable on all molecules of species α (Fig. 1c). The direction of this propulsive force has a finite temporal persistence given 1*/D*_*R*_, where *D*_*R*_ is a rotational diffusion coefficient. This describes the re-orientation timescale of the direction of actin-driven planar active force (Fig. 1c). With all the above forces, we solve the Langevin equation of motion to simulate the lateral motion of the cadherin molecules (see details in Appendix A).

### D. Recycling of cadherins

Endocytosis have been identified as an important regulatory factor for sizes of E-cadherin clusters at different surface densities in growing Drosophila embryos [33]. Multiple endocytic routes have already been identified for E-cadherin [30]. A fraction of endocytosed E-cadherins are degraded and the rest are recycled back to the membrane. Dynamin-dependent endocytosis is a major internalization process observed in Drosophila embryos [33] that selectively removes clusters from the cell surface in a size dependent manner. We incorporate this important phenomenon in our model in form of a probabilistic removal of clusters from the surface and deposition of monomers back to the surface at fixed intervals.

We implement cadherin recycling (E- or N-alike) as a stochastic removal-deposition, in spirit of the recycling kernel described by Lecuit and co-workers [33]. This kernel translates into the following removal probability for a cluster of size *c* (see Appendix A for details),

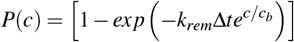

where *k*_*rem*_ is a constant removal rate describing the strength of endocytosis. The term 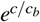 implements a removal probability that exponentially grows with the cluster size. This term ensures that any cluster of size *c* greater than the characteristic size *c*_*b*_ is almost readily removed, the removal of monomers is negligible and clusters smaller than *c*_*b*_ are removed over finite time periods. A very similar size-dependent recycling of membrane domains have been described by Rautu and coworkers in a different context [45]. Based on previous observations [33], we incorporate an exocytosis process as well in form of deposition of same number of monomers back on the surface at random locations following the removal process. This combined removal and insertion, illustrated in Fig 2a, preserve the total number of molecules at any time instant and capture the essence of endocytosis and excocytosis.

**FIG. 2.**
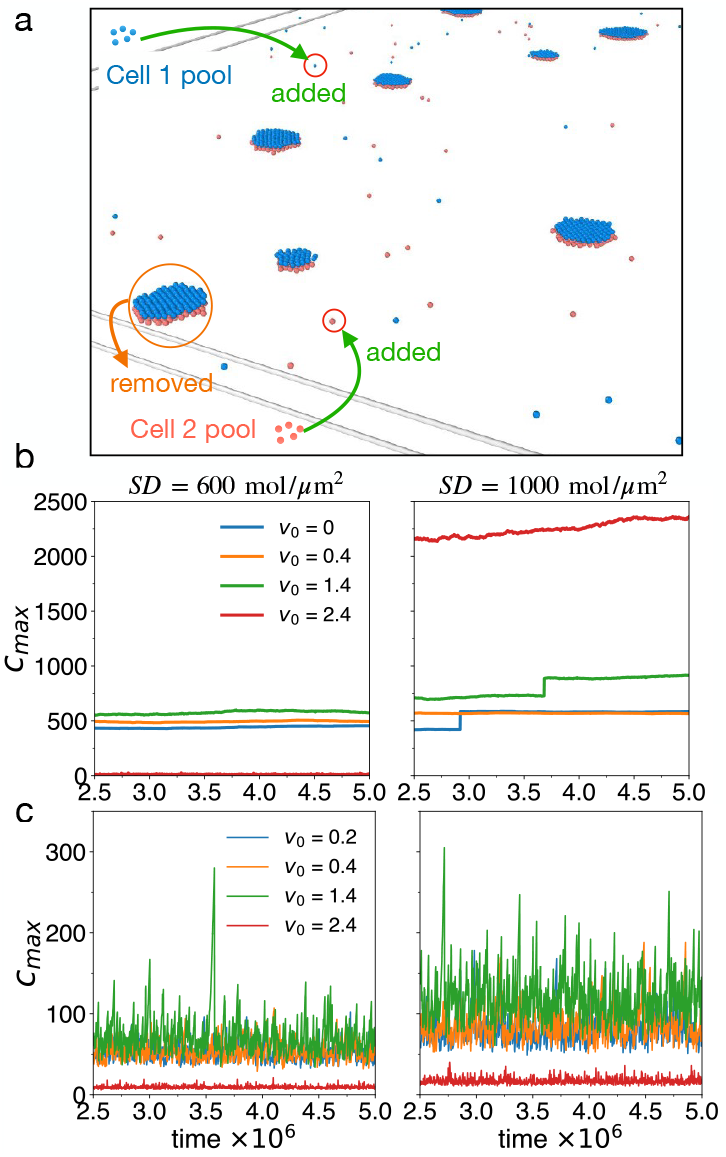
Effect of recycling on cadherin clustering. (a) Schematic of cadherin recycling in our model that shows removal of clusters from both the membranes, representing endocytosis. Monomers are added to both the cell surfaces at the same time from the virtual proteinpools of respective cells. This two steps constitute the recycling process. (b-c) Timeseries of largest cluster size *C*_*max*_for different *v*_0_ are shown at two different *SD*. Data from simulations without recycling are shown in b and with recycling and shown in c.

## III. RESULTS

### A. Fission and fusion of clusters at finite propulsive force

First we focus on the simulations with pure E-cadherin junctions in absence of recycling. Our simulations at *v*_0_ = 0 shows monotonous growth of clusters always leading to formation of several big clusters at higher surface densities (Fig. 1f). This behavior for ε_*EE*_ *>> k*_*B*_*T* is indicative of spontaneous formation of aggregates since clustering is energetically favorable. At sufficiently high densities, the clusters show gradual coalescence. Our simulations at finite *v*_0_ show a steady state with continuous occurrence of merger of small clusters along with fission of large ones. This feature recapitulates the role of actin dynamics in maintenance of clusters of E-cadherins as suggested earlier [33]. The transient propulsion helps the small clusters to find others and merge. On the other hand, several events of reorientation of the directors of propulsive forces on different molecules in a cluster leads to its disintegration into smaller ones. These two cases are highlighted in Fig. 1d and e, respectively.

When recycling is inactive, the fate of the system is decided by the interplay of activity and equilibrium clustering. At moderate densities, we observe a non-monotonic growth dynamics (Fig. 1f-top row) where clustering is suppressed at high activity. At higher densities, however, equilibrium clustering prevails over activity and we observe (Fig. 1f-bottom row) a progressive build-up of a few big clusters which eventually fuse in bigger ones.

### B. Recycling resists accumulation of big clusters and limits maximum cluster size

In absence of recycling (Fig. 2b-left), the size of the maximum cluster (*C*_*max*_) present at any instant exhibits negligible variation for a moderate *SD* and all *v*_0_. Here we express the cluster size, *c* as number of molecules in a cluster. For a higher density (Fig. 2b-right), *C*_*max*_ grows steadily in time. We observe sudden jumps in the timeseries which indicate merger-events.

When recycling is active, removal of clusters impacts the cluster formation dynamics at all densities in similar manner (Fig. 2c). The temporal dynamics of *C*_*max*_ shows significant fluctuations with intermittent growth and break-down events. The average *C*_*max*_ remains a non-monotonic function of *v*_0_ across all *SD* consistently, unlike the scenarios with inactive recycling. Clearly, the recycling mechanism, when turned on, counters the equilibrium clustering and allows the cluster maintenance via activity to continue.

### C. Recycling captures the statistical distribution of cluster-sizes observed in vivo

The statistical distribution of cadherin cluster sizes has been reported [33] to have the following functional form: 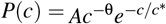 in growing Drosophila embryos. In absence of recycling, we observe *P*(*c*) to be biphasic (Supplemental material or SM figure Fig. S1) - either a power-law or exponential for *c* ≤ 10 and nearly uniform for bigger clusters. For all SDs, *P*(*c*) exhibits a significant frequency of very big clusters with *c >* 100 (Fig. S1). However, the recycling mechanism is able to prevent formation of big clusters and captures the experimentally reported form of *P*(*c*). Figure 3 shows *P*(*c*) for different *SD* and *v*_0_ combinations, and respective fitted curves and how the fitting parameters vary with *SD* and *v*_0_. We ob-serve opposite trends in the power law exponent θ (*i*.*e*. when viewed as a function of *SD* or *v*_0_ separately. The magnitude of θ decreases leading to a slower decline in cluster frequency as *SD* increases at a fixed *v*_0_ (Fig. 3a). Conversely, the magnitude of θ increases leading to faster decline in *P*(*c*) when *v*_0_ increases at a fixed *SD* (Fig. 3b). Figure 3c and d shows how different parameters of the distribution, namely *A*, exponent θ and the exponential cutoff *c*^*\**^ depend on *SD* and *v*_0_, respectively. These trends nicely capture the experimentally observed behavior and thus indicate that our simplistic model provides a viable understanding of the physical mechanism of E-cadherin clustering at the CCJs. Additional plots showing how the fitting parameters depend on the cluster removal rate *k*_*rem*_ and characteristic cutoff size *c*_*b*_ included in the recycling kernel are provided in SM figure Fig. S2.

**FIG. 3.**
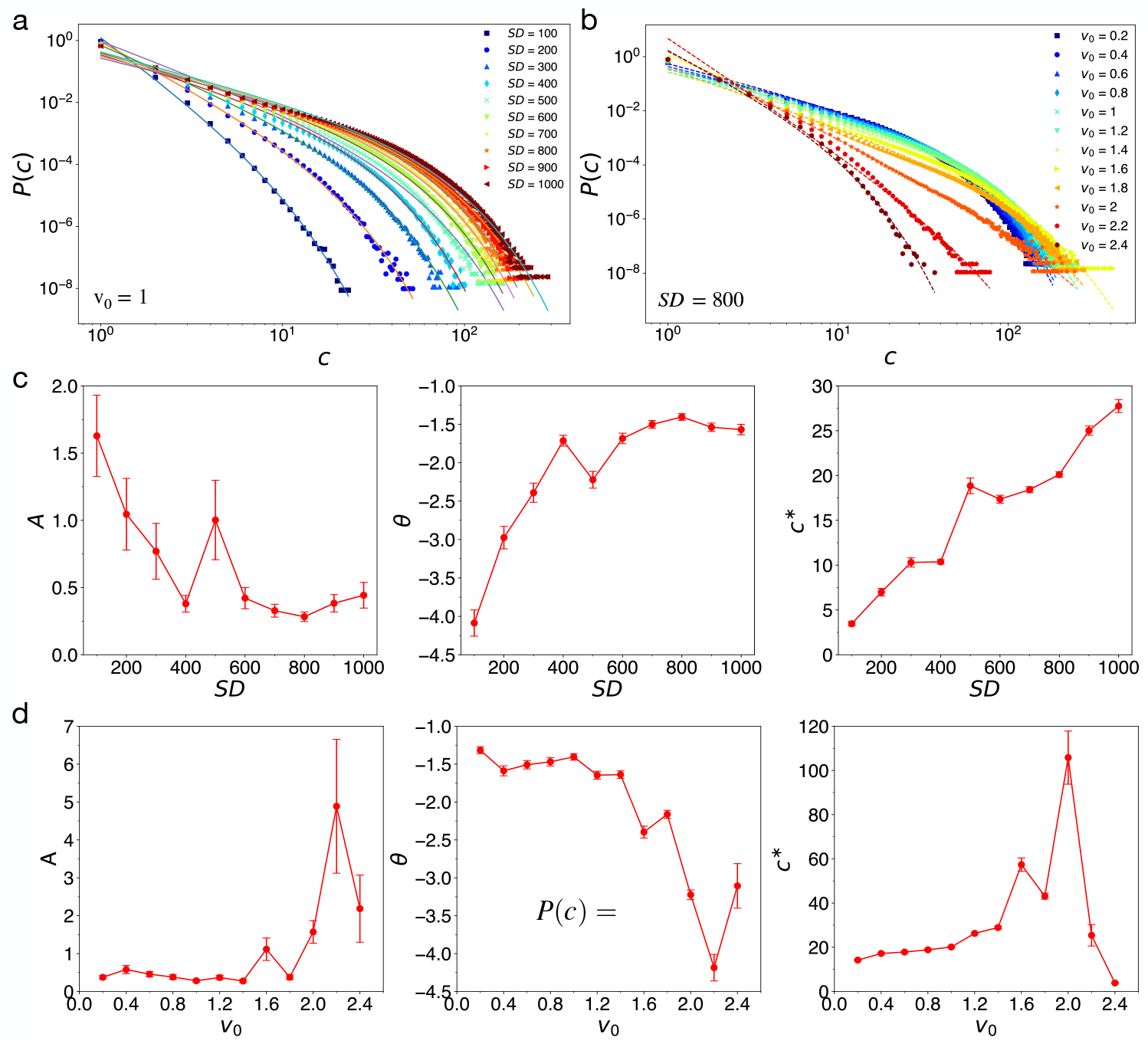
Statistical distribution of cluster size of simulated E-cadherin junctions under various conditions. Probability distribution function *P*(*c*) shown (a) for a fixed *v*_0_ = 1 at different *SD* and (b) for a fixed *SD* = 800 molecules*/µm*^2^ at different *v*_0_. The *P*(*c*) data are shown using symbols. The fits to each curve are shown using solid lines. The three fitting parameters are shown as functions of (c) *SD* and (d) *v*_0_. In both c and d, behaviors of parameters shown for *A* in left, θ in middle and *c\** in right. The errorbars in c and d are standard errors obtained from fitting. Throughout we have used *k*_*rem*_= 20 and *c*_*b*_= 1000, unless mentioned otherwise.

### D. Clustering dynamics exhibits a near-critical behavior driven by activity

Figure 3d reveals an interesting role of activity *v*_0_ in dictating the clustering dynamics. The variation of the fitted exponential cutoff parameter *c*^*\**^ shows a sharp peak around *v*_0_ = 2 which hints at emergence of a critical behavior in the clustering dynamics. We investigate this in Fig. 4. Fig. 4a shows a heatmap of *c*^*\**^ showing band of sharp peaks, which corre-sponds to very high exponential cutoff’s in these states. A large value of *c*^*\**^ indicates that *P*(*c*) here behaves nearly as a pure power-law function of 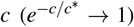. This region shows a remarkable concurrence with the cluster-depleted states (Fig. 4b). Subsequently, we discover that these states also show strong temporal fluctuations in *C*_*max*_ (Fig. 4c) with large relative variances (Fig. 4d). From a phenotypic standpoint these states may correspond to unstable junctions with a few very large E-cadherin clusters which intermittently form and breakup. Such states, however, disappear in absence of recycling of the protein molecules.

**FIG. 4.**
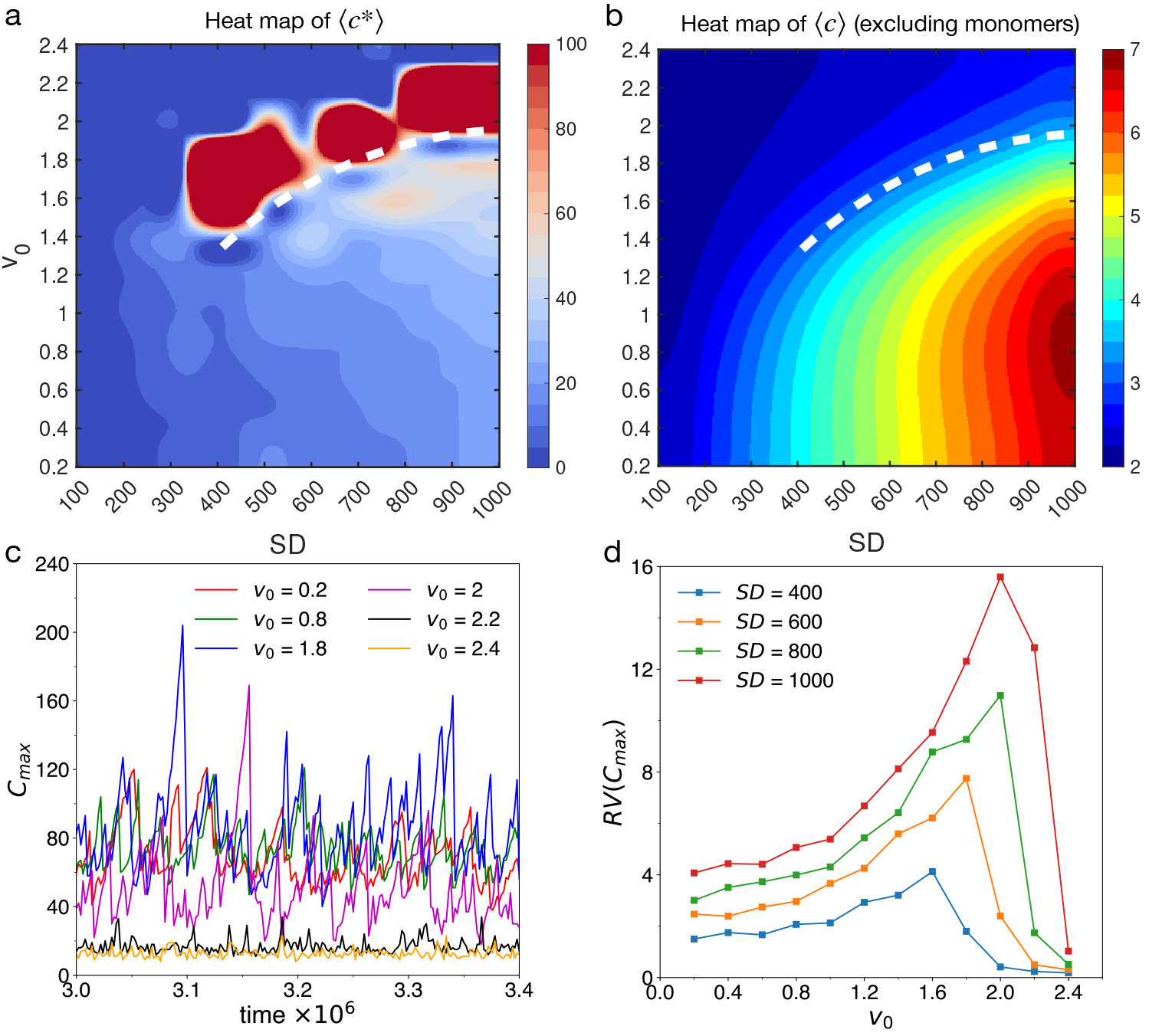
Signature of near-critical fluctuations in the clustering dynamics driven by activity and recycling. (a) Exponential cutoff *c\**, estimated from fitting *P*(*c*), shown as a function of *SD* and *v*_0_. A broken dashed line marks the boundary of a band of sharp peaks on the heatmap of *c\**. (b) Average cluster size ⟨*c*⟩ (calculated excluding the monomers and averaged over time and replicates) shown as a function of *SD* and *v*_0_, where the same broken dashed line as in panel a coincides with the boundary between states with strong and weak clustering. (c) Time-evolution of *C*_*max*_ for different *v*_0_ at *SD* = 800 molecules *µm*^2^. (d) Relative variance in *C*_*max*_, denoted as *RV* (*C*_*max*_) as a function of *v*_0_ for different *SD*.

### E. Quantification of junction strength across cadherin isoform switch

The cadherin isoform switch is known to lower the strength of epithelial junctions. Biochemical studies primarily in-fer junctional strength qualitatively, using e.g., intensities of E-cadherin immunofluorescence as proxy for junctional tension [46]. In the same spirit, we use cadherin cluster size as a measure for junction strength. We also define an interfacial adhesion energy (*U*_*adh*_), which could be a more direct reporter of junction strength. We calculate *U*_*adh*_ as the sum of all *trans* interaction energies per unit area: 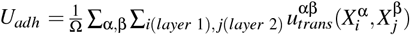. Here all species pairs included in the calculations reside on different membrane surfaces (denoted as layer 1 and 2), and Ω, the area of the junction patch, is defined in *µm*^2^.

First, we focus on cluster size as a measure of junction strength. The maturation of epithelial junctions are known to be driven by growth of clusters of E-cadherins [39]. Reduction or complete loss of E-cadherin levels at the surfaces of the adhering cells directly lowers strength of the junctions [3, 46– 51] and impacts growth of big clusters [33]. Therefore, we can expect that adherent junctions are supported by clusters of E-cadherin that are big enough in size. Motivated by this argument, we define an optimal cluster size, *c*_*opt*_. We argue that a stable junction should maintain at least a critical number, say *N*_*opt*_, of clusters with size *c* ≥ *c*_*opt*_. Evidently, *N*_*opt*_ is directly controlled by the E-cadherin levels at the cell membrane. We estimate *c*_*opt*_ at *SD* = 800 molecules*/µm*^2^, since this value represents a *SD* typical for stable, maturing junctions [33, 52]. For this, we inspect the heatmap of partially averaged cluster sizes (SM figure, Fig. S3a) that emphasizes the contribution of big clusters. We observe that this quantity shows a strong non-monotonic dependence on *v*_0_ and reaches maximum at *v*_0_ = 1.4. We choose this value as our estimate of *c*_*opt*_ ≈15 at this *SD* and *v*_0_. Next, we perform cadherin switch by replacing E-cadherins gradually with N-cadherin with its weakest *trans*-interaction, given by 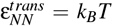 and same activity as E-cadherin. The statistical distribution *P*(*c*) for the hybrid junctions shows a preservation of the usual form, with the power-law exponent θ decreasing across the isoform switch (SM figure Fig. S3b).

Next, we estimate *N*_*opt*_ per *µm*^2^ in the model junctional area. The timeseries of *N*_*opt*_ (Fig. 5a) indicates a consistent presence of more than 10 clusters larger in size than *c*_*opt*_ per *µm*^2^ in hybrid junctions enriched in E-cadherin (*i*.*e*. low % N-cadherin). This number rapidly becomes half when only 30% switch is complete. Therefore, the hybrid junctions become significantly weaker compared to mature junctions, albeit maintaining a higher proportion of E-than N-cadherin. We observe a decrease in termporal fluctuations also as the cadherin switch takes place. The ensemble average of *N*_*opt*_ for hybrid junctions, 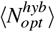 relative to the pure junction 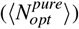, clearly portrays the above trend (Fig. 5b). It falls sharply at initial stages of cadherin switch, then reaching a saturation. The ensemble averaged interfacial adhesion energy, ⟨*U*_*adh*_⟩ for the hybrid junctions exhibits a strong correla-tion with 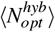, validating it as a direct measure of junctional strength. The value of ⟨*U*_*adh*_⟩ for the pure E-cadherin junction shows a good resemblance to adhesion energy values expected for *trans*-dimers of E-cadherin at the current *SD*, computed based on rupture forces measured for single E-cadherin linkages [10, 11, 53]. N-cadherin linkages show a lower rupture force [12] which is expected to lower the adhesion energy that we capture here.

**FIG. 5.**
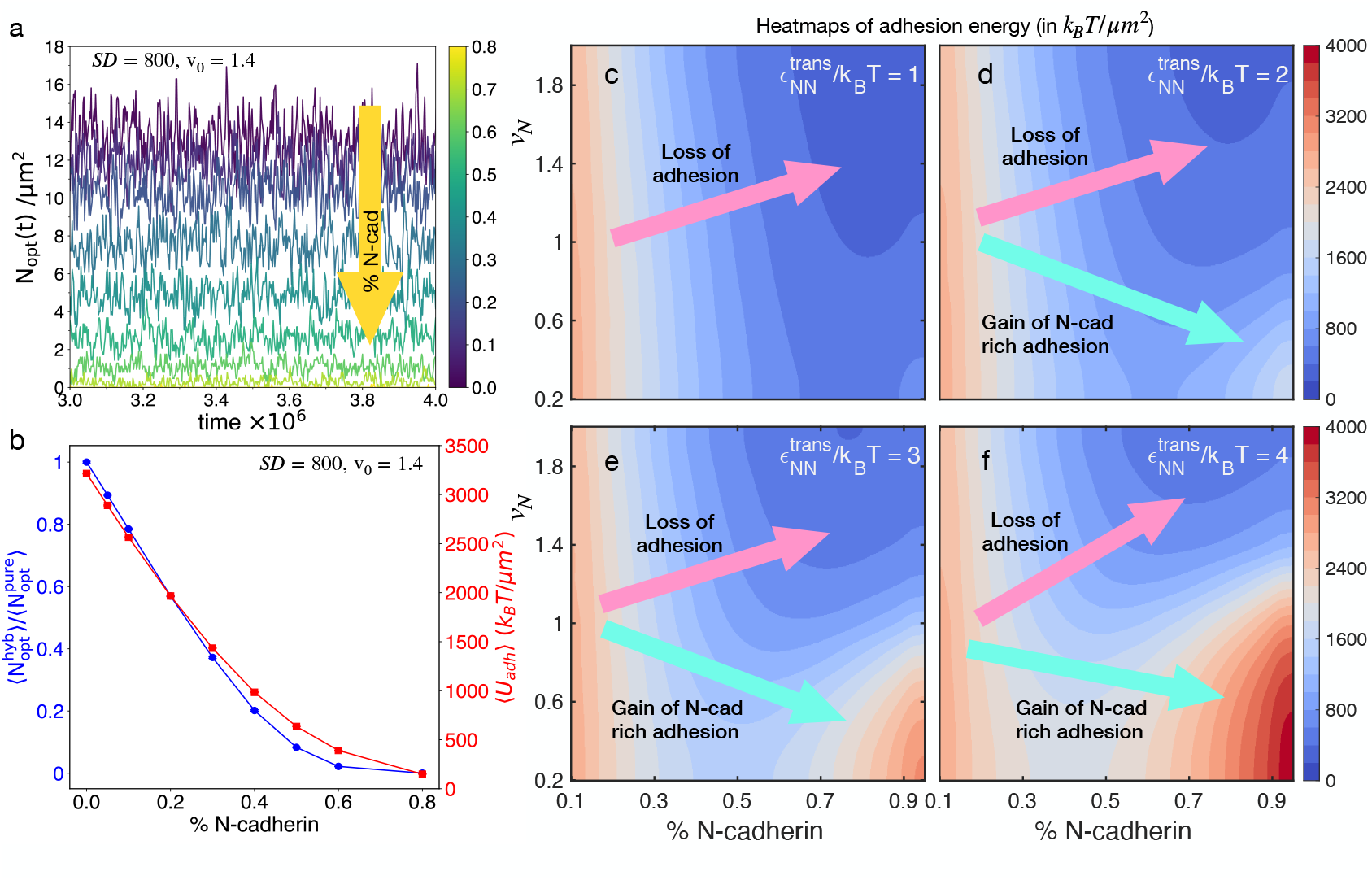
Different measures of junction strength evolve during cadherin isoform switch. (a) Timeseries of cluster count for *c* ≥ *c*_*opt*_, defined as 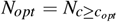, at different proportions of N-cadherin (% N-cadherin). These data highlight that the number of clusters larger than an optimal threshold gradually decreases as cadherin switch takes place. (b) Average *N*_*opt*_ for hybrid junctions relative to that in pure junctions, defined as 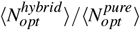 is shown to quickly decline as % N-cadherin increases. The ensemble-averaged *trans* adhesion energy ⟨*U*_*adh*_⟩ of the hybrid junctions, expressed in *k*_*B*_*T/µm*^2^, is also shown to follow the same trend with a strong correlation. In panels a and b, we have used 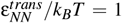. (c-f) *trans* adhesion energy as a function of % N-cadherin and the relative active propulsion of N-cadherin, defined as 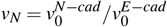. We present the heatmap of ⟨*U*_*adh*_⟩ under four different conditions: (c) 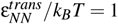, (d) 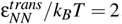, (e) 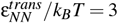, (f) 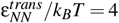. The heatmaps show gradual emergence of two behaviors at high % N-cadherin, states depicting ⟨*U*_*adh*_⟩ ≈ 0, and those with ⟨*U*_*adh*_⟩ comparable to the stable junctions with 100% E-cadherin. Possible paths leading towards these two states starting from stable adhesion states (*v*_*N*_ ≈ 1) are shown using solid arrows of colors purple and cyan, respectively, as guide in all the panels.

### F. A bi-modal behavior of actively driven junctions emerges from the cadherin isoform switch

Now we focus on how the strength of hybrid junctions are influenced by the interplay of activity and energetics of Eand N-cadherin *trans*-linkages. This exercise would lead us to different junctional phenotypes. We inspect the behavior of ⟨*U*_*adh*_⟩ as we tune the activity and % N-cadherin. Ncadherins have been known to interact with cortical actin differently compared to E-cadherin and hence offers a weaker but more dynamic adhesive capability [10–12, 54, 55]. We capture this behavior of N-cadherin within our model by tuning the active propulsion for N-cadherin relative to E-cadherin. Thus we define a relative activity for N-cadherin, given by 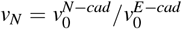 which we tune in conjunction with the cadherin isoform switch. We also vary the strengths of transinteraction for N-cadherin from very weak (*k*_*B*_*T*) to as strong as E-cadherin (4*k*_*B*_*T*).

Figure 5c-f shows the changes in adhesion energy (⟨*U*_*adh*_⟩) of the hybrid junctions as a joint function of *v*_*N*_ and % N-cadherin. We observe that for very weak N-cadherin *trans*-contacts (Fig. 5c-d), ⟨*U*_*adh*_⟩ remains low for the entire spectrum of cadherin switch. We see emergence of two regions of adhesion at high % N-cadherin, as the N-cadherin *trans*-interaction strength increases (Fig. 5e-f): states with low adhesion energy (⟨*U*_*adh*_⟩ ≈ 0) and states with high adhesion energy (⟨*U*_*adh*_⟩ *>* 3000 *k*_*B*_*T/µm*^2^, which is similar to the pure E-cadherin condition, depicted in Fig. 5b. The low adhesion energy states always emerge at high value of *v*_*N*_ which reflects the experimentally implicated scenario of N-cadherin being driven by highly dynamic cortical actin [12, 55]. This situation would also indicate that the adhesion is weak when N-cadherin clusters remodel rapidly. Such a condition would be helpful for maintaining weak and dynamic adhesions at the cell-cell interface that is necessary for fast, collective migration. This behavior has been seen recently in cancer-associated fibroblasts [54]. The high adhesion energy states always emerge at small *v*_*N*_ which means in such cases N-cadherin is weakly driven by F-actin. This would mean that the junctions are less dynamic and stabilizing via long-lived N-cadherin mediated adhesions as observed in neural tissues [56]. Both these two types of states are potential junctional phenotypes in the hybrid-E/M spectrum. The heatmaps in Fig. 5c-f collectively represents a simplified landscape of the hybrid E/M-like junctional phenotypes.

### IV. DISCUSSION AND CONCLUSION

In this work, we have developed a physical model to decode the non-equilibrium features of cadherin-based CCJs in biological tissues. Here we model the E- and N-cadherin extracellular domains as spherical molecules to predict a min-imal energy landscape of diverse junctional states. We consider the most fundamental features of cadherin dynamics at the cell membrane: lateral diffusion of cadherin extracellular domains, transient propulsion of the cadherin molecules inflicted by cortical actin and myosin, and an exponential recycling process. The striking flexibility and robustness of the model is revealed when we could adapt the same framework to capture the junctional statistical distribution of E-cadherin clusters in Drosophila embryos [33] as well as junctional features of hybrid E/M cells.

Subsequently, we identify the conditions for epithelial-like, stable CCJs, and perform cadherin switching by gradually replacing E-cadherin molecules with N-cadherin. In the model, we differentiate these two homologous species by two factors based on literature: the intermolecular adhesion strength and the magnitude of actin-driven propulsive force. We compute the junctional adhesion energy for low to high adhesion strengths and small to large magnitudes of propulsive forces to predict diverse junctional phenotypes. These phenotypes include stable junctions (high adhesion energy) enriched in E-cadherin, or N-cadherin and weak hybrid junctions (low adhesion energy) containing a small to large proportion of N-cadherin. Based on information about cadherin populations in CCJs, we can map the pure E-cadherin-based junctions to epithelial junctions. On the other hand, junctions with only N-cadherins are observed with both low and high adhesion energy, which may be mapped to the junctions in mesenchymal cells with abilities to form a range of adhesions of variable strengths. Any junction with both varieties of cadherins are found to have low to intermediate strength and can be mapped to the junctions in hybrid E/M cells. These states are schematically shown in a 3D version of the minimal EMT landscape in Fig. 6, along with one of the many possible pathways for EMT. Such a landscape provides the foundations for understanding cell state switching processes driven by mechanochemical coupling in many other biological contexts.

**FIG. 6.**
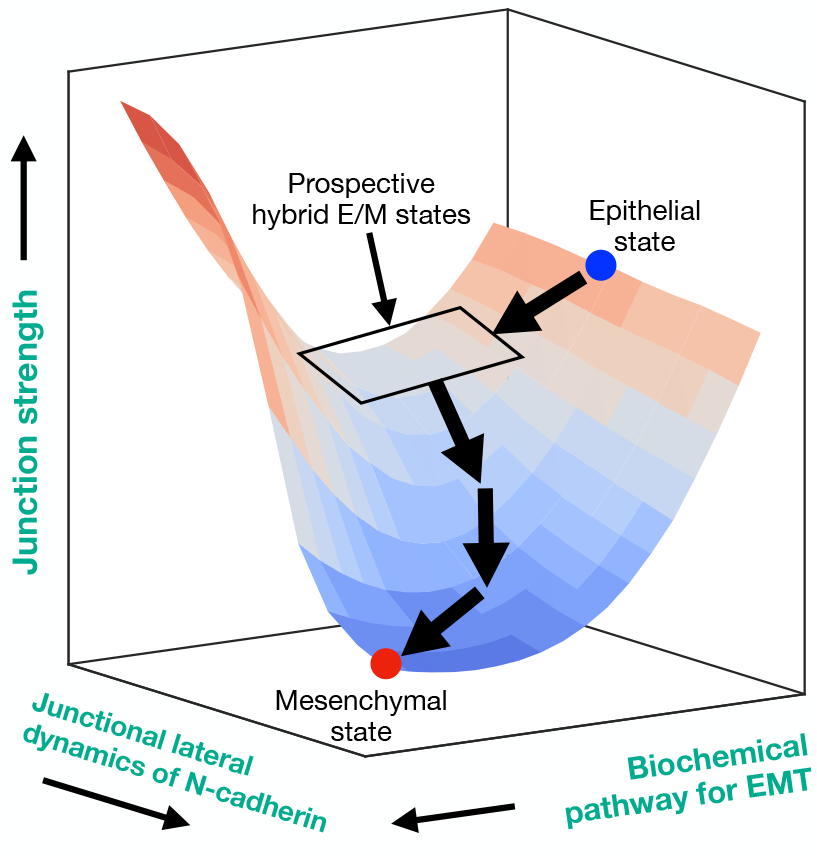
A minimal landscape of EMT, highlighting the hybrid E/M states and possible paths for EMT. Here, junction strength is represented by the measured adhesion energy, lateral dynamics of N-cadherin by *v*_*N*_ and biochemical pathway for EMT by % of N-cadherin. This landscape is a representation for how mechanochemical coupling dictates the process of EMT. The hybrid E/M states emerge as a valley in this landscape where the junction strength is intermediate to the terminal states. Because of the proximity of these states to the epithelial state on the mechanochemical plane, they can revert back and initiate secondary tumors. Junction strength in this region is intermediate and remains quite similar across these states. These features allow them to modulate the junctional configurations with minimal cost and gives them plasticity.

All the junctional phenotypes we observe here have been either observed earlier or have been implicated in specific situations - but never connected through a consistent measure of junction strength and pin-pointed on a map of states. Pure E-cadherin *trans* contacts get more stabilized when pulled by cortical contractile forces from F-actin via the catch-bond effect [32]. This helps the junctions to mature and reinforce. This behavior makes E-cadherin the necessary adhesion receptor in epithelial tissues. N-cadherin, on the other hand, is much less effective at reinforcement and rather responsive to dynamic actin reorganization. Recent evidence show that the N-cadherin *trans*-dimers may have a larger unbinding rate from actin compared to E-cadherin [12]. There is significant evidence that N-cadherin *trans*-contacts break down at nearly an order of magnitude much lesser extent of force compared to E-cadherin [10, 11]. This makes N-cadherin *trans*-contacts energetically less stable and more prone to dynamic organization. In the context of EMT, this behavior is relevant for the mesenchymal cells which are more susceptible to maintain weak and dynamic cell-cell adhesions. Mesenchymal cells dedicate majority of their actin machinery to migrate effectively. This behavior has been seen recently in cancer-associated fibroblasts where N-cadherin plays a major role helping migration, maintaining weaker adhesions at the cellcell interfaces and stronger adhesions at the cell-ECM interface [54]. Another recent work has pointed out N-cadherin’s role at maintaining a high cortical actin activity via controlling RAC1-signaling [55] which also support these observations.

The high adhesion energy states observed for nearly pure N-cadherin junctions at low active propulsion could be an interesting phenotype to investigate in future, possibly through appropriately designed experiments. A possible design might include an *in-vitro* reconstituted cortex on a supported bilayer where actomyosin activity and N-cadherin concentration can be tuned. If a second layer of N-cadherin molecules, all fixed on a functionalized surface is brought in contact with the actively driven layer of N-cadherin - we might be able to study the effects of active propulsion on the N-cadherin junction strength in a controlled setting. In cellular context, neural progenitor cells or mesenchymal stromal cells found in connective tissues around tumors may be candidates to study similar junctional phenotypes. can Actomyosin activity in cells maybe controlled by introducing appropriate drugs or a mutated isoform of myosin with lowered activity.

From a physics point of view, the most striking feature of the model is the interplay between active propulsion and recycling. The latter turns out to an essential ingredient in the current model that not only works as a pathway to removal of large cadherin clusters but also acts as a different type of non-equilibrium driving. In context of clustering of colloidal particles of few nanometer size, effects of active propulsion is well studied [35, 57]. Size-dependent recycling has been used to describe microphase separation of lipidic membranes [45]. However, an investigation into an interplay of these two factors is fairly unexplored till this point to the best of our knowledge. Our model brings out the steady state features of a clustering mechanism driven by propulsion and recycling and reveals the principles of cluster maintenance at a mesoscopic scale. Similar in spirit to the fluctuation-dominated ordering shown previously [43], we pinpoint a strongly nonequilibrium region on the density-activity plane where effects of recycling seem to get amplified by the active driving provided by the molecular level propulsion. A possible mechanism could be the following: optimal values of propulsion leads to maximal cluster-fusions and as a result very large clusters are formed which are removed at a very high rate. This way formation and break-up dynamics are magnified under these conditions.

Finally, we also highlight how these essential biophysical cellular mechanisms, can also drive phenotypic shifts in CCJs made of two kinds of adhesion receptors with differential coupling to cortical active forces. In principle, they could also have differential recycling behaviors as well. We could not explore that here because of limited literature available on recycling mechanisms of N-cadherin. We expect that any differences in the endo- or exo-cytic pathways of the two cadherins could potentially lead to variations in the junction strengths and produce a richer hybrid E/M landscape. We plan to pursue this avenue through a future effort in this direction.

## APPENDIX A

### A. Details of the simulations

We simulate the lateral motion of the cadherin molecules on the membrane via solving the Langevin equation of motion, which for the *i*th molecule of the species α is given by,

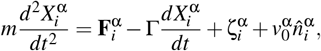

where *m* is the mass of any molecule assumed to be uniform across species. 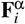 is the total force on *i*th molecule of the species α due to all pairwise interactions. 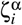 is the Brow-nian random force which obeys the following: ⟨ ζ(*t*)⟩ = 0 and ⟨ζ(*t*)ζ(*t*′)⟩ = 2*dk*_*B*_*T* Γδ(*t*− *t*′) where *d* = 2, the dimensionality of the motion. The temporal Dirac delta function, δ(*t* − *t*′) = 1*/*Δ*t* with Δ*t* being the discrete timestep of our simulation. The magnitude of the random force is chosen via the fluctuation-dissipation theorem to be consistent with the specified drag and temperature.

The term 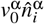 describes the lateral propulsive force expe-rienced by the *i*th molecule of species α where 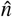 depicts the direction of the force on the flat surface: 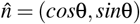. The angle θ (illustrated in Fig. 1c) evolves in time as: 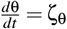. The quantity ζ_θ_ is a Gaussian white noise with zero mean and following correlations: ⟨ζ_θ_(*t*)ζ_θ_(*t*′⟩ = 2*D*_*R*_δ(*t* − *t*′).

We use the python-based open source package HOOMD-Blue [58] to implement our model and perform all the simulations. We implement the above lateral dynamics and recycling process for two adjacent membrane surfaces held together at a separation of σ. We consider different surface densities (SDs) of cadherins in the range 100 − 1000 molecules per *µm*^2^. We capture these densities with a fixed 5000 molecules per membrane surface and varying the non-dimensionalized lateral dimension *L* of the simulated region in the range 833σ − 417σ. Thus, we simulate the dynamics of 10000 molecules in total. All pair interactions among adjacent molecules are computed with a cutoff radius of 2.5σ. So, molecule-pairs found at a separation more that this value do not interact. We vary the strength of the propulsive force *v*_0_ between nondimensionalized values [0 − 3] with fixed *D*_*R*_ = 1. Such propulsive forces range between 0.1 to 1 *pN* which is similar to those observed for single myosin motors exerted on actin filaments [59]. In majority of our calculations, the exponential recycling is performed with a constant removal rate, *k*_*rem*_ = 20 molecules per 1000 time steps and *c*_*b*_ = 1000, unless mentioned otherwise. In SM figure S2, we have presented effects of varying *k*_*rem*_ in range [10 − 50], and *c*_*b*_ in range [100 − 1000].

We integrate Langevin equations of motion for the cadherin molecules with a timestep Δ*t* = 0.01 in simulation time unit. Each of our simulations are 5 × 10^6^ steps long. We generate 10-20 replicates for different combinations of control parameters (*v*_0_, *SD, c*_*b*_, *& k*_*rem*_) to analyze the probability distribution of cluster sizes.

We express all energies in the model in terms of *k*_*B*_*T* with temperature *T* set at the physiological 310K. The lengths in the model are expressed in terms of the average crosssectional diameter of any cadherin molecule (E or N) which has been reported ∼ 6 *nm* [33]. We fix the time unit τ = 10 *ps* based on a suitable value of the ratio *m/*Γ that corresponds to a lateral diffusion coefficient of ≈ 1 *µm*^2^ *s*^*−*1^ for the single cadherin molecules as observed previously in literature for similar transmembrane proteins [60].

### B. Detection of clusters

To detect the clusters in our simulated configurations, we use the python module - freud [61] that detects clusters based on nearest neighbors, which are particles located within a defined cutoff distance (*r*_*max*_). For each frame of the simulation, freud computes the proximity of particles based on their positions in the simulation box and the cutoff *r*_*max*_. We set *r*_*max*_ = 1.5 to detect the clusters. The algorithm also takes care of the periodic boundary conditions and ensure clusters are correctly identified even when particles are near the boundaries of the simulation area. Once clusters are identified, each particle is assigned a “cluster index,” which indicates to which cluster the particle belongs. These indices allow for tracking and analyzing the structure and dynamics of individual clusters. After detection of the clusters, the freud package computes the size of each cluster, defined as the number of particles within a cluster. This allows us to compute the statistical distribution of the cluster sizes.

### C. Derivation of removal probability for clusters of given size

As described in Ref [33], the exponential cluster recycling process can be represented by the following differential equation:

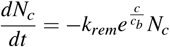

where *N*_*c*_ is the number of clusters of size *c* present at time *t* per unit surface area. *k*_*rem*_ is the rate constant for removal of clusters, describing how quickly clusters are being removed. *c*_*b*_ represents a characteristic cutoff cluster size which controls the exponential growth of the removal rate with cluster size *c*. This equation describes a process where clusters of various sizes are subjected to a selective depletion. Specifically, we observe that small clusters (*c << c*_*b*_) get depleted slowly be-cause 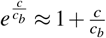. Large clusters (*c >> c*_*b*_) get depleted even more quickly at exponentially amplified rates. Solution to this equation is: 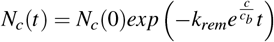. This means the number of a certain cluster size decays exponentially in time, and interestingly the rate of decay itself is exponentially increasing with cluster size. Here we implement the solution as a Monte-Carlo module to simulate the removal process in a stochastic manner. We do this by randomly removing any cluster with a depletion probability. From time 0 to Δ*t* the number of clusters of size *c* changes from *N*_*c*_(0) to *N*_*c*_(Δ*t*). The depletion probability would then can be written as:

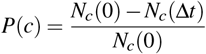

Using the solution above we can write:

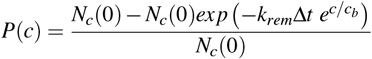

Rearranging we get 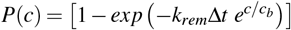 as mentioned in the main text.

## ACKNOWLEDGEMENTS

Authors thank Anjan Roy, Ishaan Gupta and members of AD laboratory for helpful discussions. KS acknowledges IIT Delhi for a fellowship. AD acknowledges funding support from the New Faculty SEED Grant from IIT Delhi, India and the Start-up Research Grant from SERB-DST, Government of India (Grant No. SRG/2023/000099). Authors acknowledge support from IIT Delhi High Performance Computing facilities.

## Data Availability

All data generated through the model are included in the article and/or supplemental material.

## References

[1] B. Alberts, R. Heald, A. Johnson, D. Morgan, M. Raff, K. Roberts, and P. Walter, Molecular biology of the cell: seventh international student edition with registration card (WW Norton & Company, 2022).

[2] M. S. Adil, S. P. Narayanan, and P. R. Somanath, Tissue barriers 9, 1848212 (2021), place: United States.

[3] S. Curran, C. Strandkvist, J. Bathmann, M. d. Gennes, A. Kabla, G. Salbreux, and B. Baum, Developmental Cell 43, 480 (2017).

[4] R. J. Huebner, A. N. Malmi-Kakkada, S. Sarıkaya, S. Weng, D. Thirumalai, and J. B. Wallingford, eLife 10, e65390 (2021).

[5] M. Nieto, R.-J. Huang, R. Jackson, and J. Thiery, Cell 166, 21 (2016), number: 1.

[6] K. K. Youssef and M. A. Nieto, Nature Reviews Molecular Cell Biology 25, 720 (2024), number: 9.

[7] C. E. Aban, A. Lombardi, G. Neiman, M. C. Biani, A. La Greca, A. Waisman, L. N. Moro, G. Sevlever, S. Miriuka, and C. Luzzani, Scientific Reports 11, 2048 (2021), number: 1.

[8] C.-Y. Loh, J. Chai, T. Tang, W. Wong, G. Sethi, M. Shanmugam, P. Chong, and C. Looi, Cells 8, 1118 (2019), number: 10.

[9] O. J. Harrison, X. Jin, S. Hong, F. Bahna, G. Ahlsen, J. Brasch, Y. Wu, J. Vendome, K. Felsovalyi, C. M. Hampton, R. B. Troyanovsky, A. Ben-Shaul, J. Frank, S. M. Troyanovsky, L. Shapiro, and B. Honig, Structure 19, 244 (2011), publisher: Elsevier BV.

[10] Y.-S. Chu, W. A. Thomas, O. Eder, F. Pincet, E. Perez, J. P. Thiery, and S. Dufour, The Journal of Cell Biology 167, 1183 (2004), number: 6.

[11] S. Huveneers and J. de Rooij, Journal of Cell Science 126, 403 (2013).

[12] H. Zhu, X. Liu, J. Zhang, G. Zhao, J. Wang, H. Zhang, Y. Liu, H. Guo, J. Yang, Z. Wang, T. J. Lu, F. Xu, and M. Lin, Biophysical Journal 124, 2041 (2025), publisher: Elsevier.

[13] N. Meyer-Schaller, M. Cardner, M. Diepenbruck, M. Saxena, S. Tiede, F. Lüönd, R. Ivanek, N. Beerenwinkel, and G. Christofori, Developmental Cell 48, 539 (2019), number: 4.

[14] M. K. Jolly, S. A. Mani, and H. Levine, Biochimica et Biophysica Acta (BBA) - Reviews on Cancer 1870, 151 (2018), number: 2.

[15] A. Grosse-Wilde, A. Fouquier d’Hérouël, E. McIntosh, G. Ertaylan, A. Skupin, R. E. Kuestner, A. Del Sol, K.-A. Walters, and S. Huang, PLOS ONE 10, e0126522 (2015), number: 5.

[16] M. K. Jolly, B. Huang, M. Lu, S. A. Mani, H. Levine, and E. Ben-Jacob, Journal of The Royal Society Interface 11, 20140962 (2014), number: 101.

[17] B. Bierie, S. E. Pierce, C. Kroeger, D. G. Stover, D. R. Pattabiraman, P. Thiru, J. Liu Donaher, F. Reinhardt, C. L. Chaffer, Z. Keckesova, and R. A. Weinberg, Proceedings of the National Academy of Sciences 114, 10.1073/pnas.1618298114 (2017), number: 12.

[18] C. Kröger, A. Afeyan, J. Mraz, E. N. Eaton, F. Reinhardt, Y. L. Khodor, P. Thiru, B. Bierie, X. Ye, C. B. Burge, and R. A. Weinberg, Proceedings of the National Academy of Sciences 116, 7353 (2019), number: 15.

[19] C. Elie-Caille, I. Lascombe, A. Péchery, H. Bittard, and S. Fauconnet, Molecular and Cellular Biochemistry 471, 113 (2020), number: 1-2.

[20] M. K. Jolly, J. A. Somarelli, M. Sheth, A. Biddle, S. C. Tripathi, A. J. Armstrong, S. M. Hanash, S. A. Bapat, A. Rangarajan, and H. Levine, Pharmacology & Therapeutics 194, 161 (2019).

[21] S. Tripathi, J. Xing, H. Levine, and M. K. Jolly, in The Epithelial-to Mesenchymal Transition: Methods and Protocols, edited by K. Campbell and E. Theveneau (Springer US, New York, NY, 2021) pp. 385–413.

[22] S. Muralidharan, S. Sahoo, A. Saha, S. Chandran, S. S. Majumdar, S. Mandal, H. Levine, and M. K. Jolly, Biomolecules 12, 10.3390/biom12020297 (2022).

[23] K. Hari, S. Tripathi, V. Anand, M. K. Jolly, and H. Levine, bioRxiv 10.1101/2024.11.26.625479 (2025), publisher: Cold Spring Harbor Laboratory.

[24] S. E. Leggett, J. Y. Sim, J. E. Rubins, Z. J. Neronha, E. K. Williams, and I. Y. Wong, Integrative biology : quantitative biosciences from nano to macro 8, 1133 (2016), place: England.

[25] J. A. Mitchel, A. Das, M. J. O’Sullivan, I. T. Stancil, S. J. DeCamp, S. Koehler, O. H. Ocavña, J. P. Butler, J. J. Fredberg, M. A. Nieto, D. Bi, and J.-A. Park, Nature Communications 11, 5053 (2020).

[26] H. Wang, R. Liu, Y. Yu, H. Xue, R. Shen, Y. Zhang, and J. Ding, Biomaterials 317, 123013 (2025).

[27] K. Hosseini, A. Frenzel, and E. Fischer-Friedrich, Biophysical Journal 120, 3516 (2021).

[28] A. Datta, S. Deng, V. Gopal, K. C.-H. Yap, C. E. Halim, M. L. Lye, M. S. Ong, T. Z. Tan, G. Sethi, S. C. Hooi, A. P. Kumar, and C. T. Yap, Cancers 13, 10.3390/cancers13081882 (2021), place: Switzerland.

[29] Y. Chen, J. Brasch, O. J. Harrison, and T. C. Bidone, Biophysical Journal 120, 4944 (2021), publisher: Elsevier.

[30] I. Woichansky, C. A. Beretta, N. Berns, and V. Riechmann, Nature communications 7, 10834 (2016), publisher: Nature Publishing Group UK London.

[31] I. Pastushenko, A. Brisebarre, A. Sifrim, M. Fioramonti, T. Revenco, S. Boumahdi, A. Van Keymeulen, D. Brown, Moers, S. Lemaire, S. De Clercq, E. Minguijón, C. Balsat, Y. Sokolow, C. Dubois, F. De Cock, S. Scozzaro, F. Sopena, A. Lanas, N. D’Haene, I. Salmon, J.-C. Marine, T. Voet, P. A. Sotiropoulou, and C. Blanpain, Nature 556, 463 (2018), number: 7702.

[32] C. D. Buckley, J. Tan, K. L. Anderson, D. Hanein, N. Volkmann, W. I. Weis, W. J. Nelson, and A. R. Dunn, Science 346, 1254211 (2014), publisher: American Association for the Advancement of Science.

[33] B.-A. Truong Quang, M. Mani, O. Markova, T. Lecuit, and P.-F. Lenne, Current Biology 23, 2197 (2013), number: 22.

[34] Y. Wu, X. Jin, O. Harrison, L. Shapiro, B. H. Honig, and A. Ben-Shaul, Proceedings of the National Academy of Sciences 107, 17592 (2010), number: 41.

[35] Y. Fily and M. C. Marchetti, Physical Review Letters 108, 235702 (2012).

[36] Y. Wu, B. Honig, and A. Ben-Shaul, Biophysical Journal 104, 1221 (2013), number: 6.

[37] Q. Yu, T. Kim, and V. Rajagopal, PLOS Computational Biology 18, e1010257 (2022), number: 7.

[38] J. Brasch, O. J. Harrison, B. Honig, and L. Shapiro, Trends in Cell Biology 22, 299 (2012), number: 6.

[39] A. S. Yap, G. A. Gomez, and R. G. Parton, Developmental Cell 35, 12 (2015).

[40] J. Chen, Z.-R. Xie, and Y. Wu, Molecular BioSystems 12, 205 (2016), number: 1.

[41] P. Katsamba, K. Carroll, G. Ahlsen, F. Bahna, J. Vendome, S. Posy, M. Rajebhosale, S. Price, T. M. Jessell, A. Ben-Shaul, L. Shapiro, and B. H. Honig, Proceedings of the National Academy of Sciences 106, 11594 (2009), number: 28.

[42] K. Gowrishankar, S. Ghosh, S. Saha, C. Rumamol, S. Mayor, and M. Rao, Cell 149, 1353 (2012).

[43] A. Das, A. Polley, and M. Rao, Physical Review Letters 116, 068306 (2016).

[44] S. Saha, A. Das, C. Patra, A. A. Anilkumar, P. Sil, S. Mayor, and M. Rao, Proceedings of the National Academy of Sciences 119, e2123056119 (2022), publisher: National Acad Sciences.

[45] S. A. Rautu, G. Rowlands, and M. S. Turner, Europhysics Letters 121, 58004 (2018), publisher: EDP Sciences, IOP Publishing and Società Italiana di Fisica.

[46] G. R. Kale, X. Yang, J.-M. Philippe, M. Mani, P.-F. Lenne, and T. Lecuit, Nature Communications 9, 5021 (2018).

[47] C. Bertet, L. Sulak, and T. Lecuit, Nature 429, 667 (2004).

[48] T. E. Vanderleest, C. M. Smits, Y. Xie, C. E. Jewett, J. T. Blankenship, and D. Loerke, eLife 7, 34586 (2018).

[49] G. Charras and A. S. Yap, Current Biology 28, R445 (2018).

[50] J. X. H. Li, V. W. Tang, and W. M. Brieher, Proceedings of the National Academy of Sciences, USA 117, 432 (2020).

[51] E. Heller, K. V. Kumar, S. W. Grill, and E. Fuchs, Developmental Cell 28, 617 (2014).

[52] S. Togo, K. Sato, R. Kawamura, N. Kobayashi, M. Noiri, S. Nakabayashi, Y. Teramura, and H. Y. Yoshikawa, APL Bioengineering 4, 016103 (2020).

[53] D. Leckband and J. de Rooij, Annual Review of Cell and Developmental Biology 30, 291 (2014), publisher: Annual Reviews Type: Journal Article.

[54] A. Labernadie, T. Kato, A. Brugués, X. Serra-Picamal, S. Derzsi, E. Arwert, A. Weston, V. González-Tarragó, A. Elosegui-Artola, L. Albertazzi, J. Alcaraz, P. Roca-Cusachs, E. Sahai, and X. Trepat, Nature Cell Biology 19, 224 (2017).

[55] S. Moazzeni, K. Kyker-Snowman, R. I. Cohen, H. Wang, R. Li, D. I. Shreiber, J. D. Zahn, Z. Shi, and H. Lin, Scientific Reports 15, 4296 (2025).

[56] Y. Miyamoto, F. Sakane, and K. Hashimoto, Cell Adhesion & Migration 9, 183 (2015), publisher: Taylor & Francis.

[57] E. Mani and H. Löwen, Physical Review E 92, 032301 (2015), publisher: American Physical Society.

[58] J. A. Anderson, J. Glaser, and S. C. Glotzer, Computational Materials Science 173, 109363 (2020).

[59] R. Phillips, J. Kondev, J. Theriot, and H. Garcia, Physical biology of the cell (Garland Science, 2012).

[60] S. Saha, I.-H. Lee, A. Polley, J. T. Groves, M. Rao, and S. Mayor, Molecular biology of the cell 26, 4033 (2015), publisher: The American Society for Cell Biology.

[61] V. Ramasubramani, B. D. Dice, E. S. Harper, M. P. Spellings, J. A. Anderson, and S. C. Glotzer, Computer Physics Communications 254, 107275 (2020).

