## Supplemental figures for "Dynamics of single cell-cell junctions as an indicator of cell state switch"

### Supplemental material for 'Dynamics of single cell-cell junctions as an indicator of cell state switch'

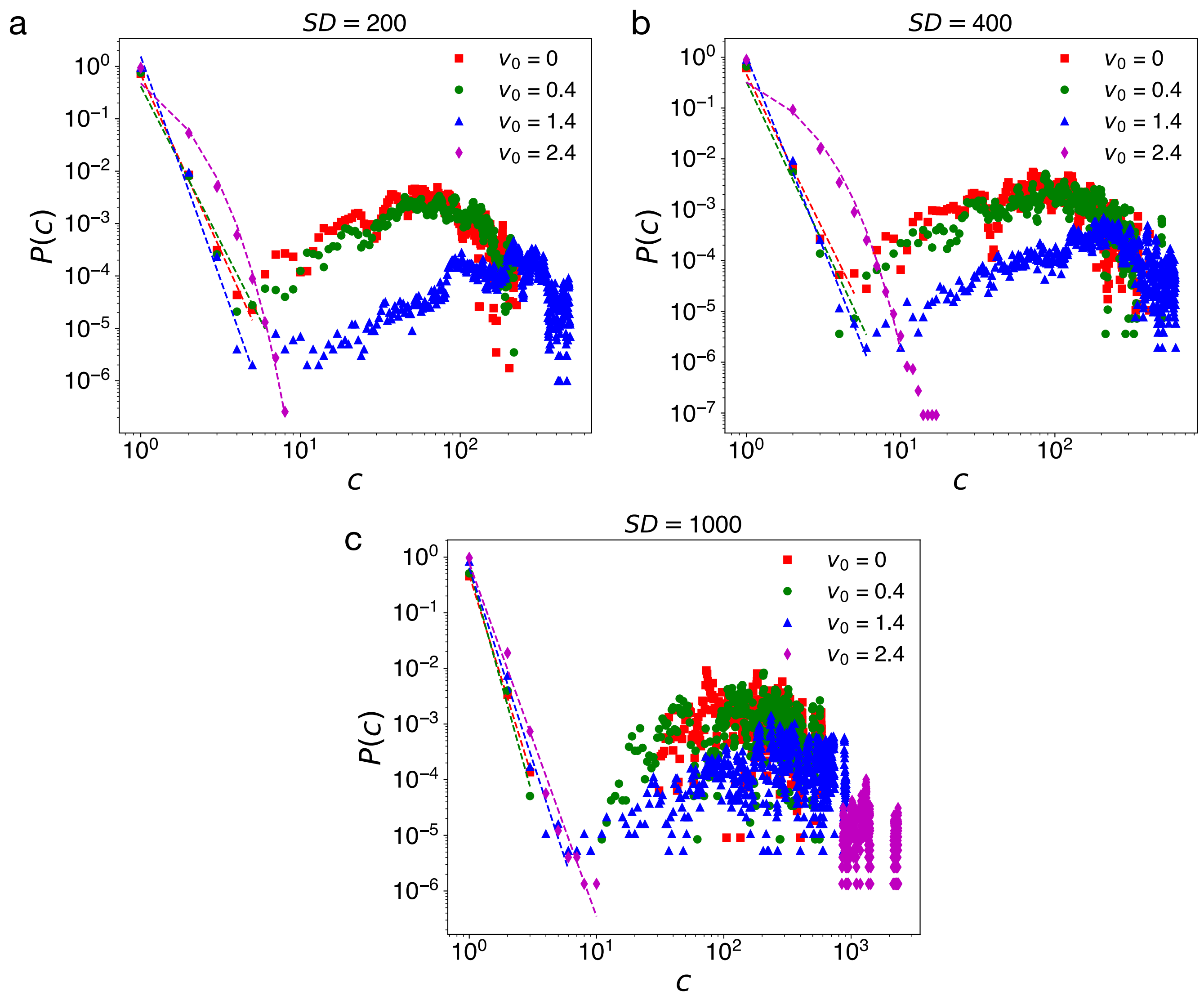


FIG. 1. Probability distribution function $P(c)$ of cluster sizes $c$ at three different surface density values: (a) $SD = 200$, (b) $SD = 400$, and $SD = 1000$ molecules/$\mu m^2$. For each case, profiles of $P(c)$ are shown at different $v_0$ values. The dashed lines shown in each panel are either power-law fits (straight lines in a, b and c) or exponential fits (curved lines in a and b). In all these cases recycling is kept inactive.

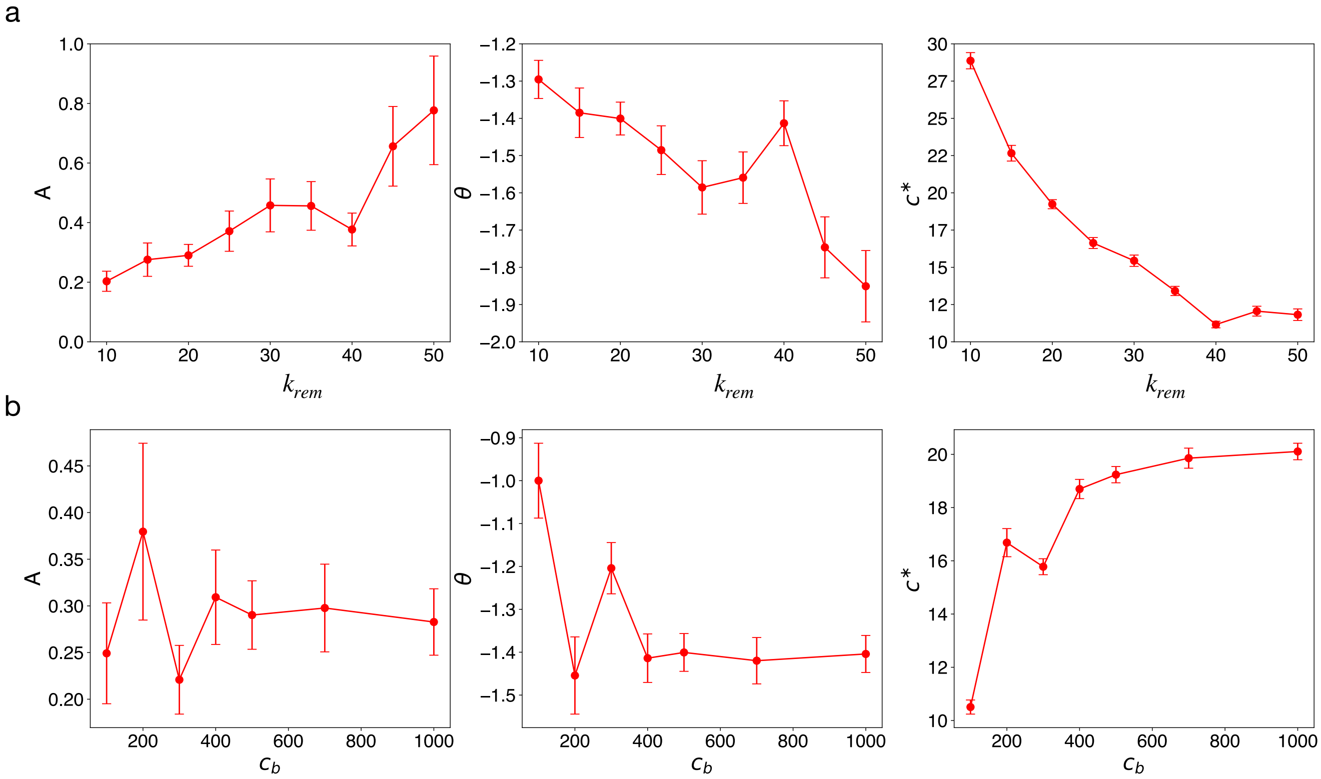


FIG. 2. Data showing how the fitting parameters of $P(c)$ depend on (a) the cluster removal rate constant $k_{rem}$, and (b) cutoff parameter $c_b$ used in the exponential part of the removal rate. In both rows from left, first panels show the values of pre-factor $A$, second panels show the power-law exponent $\theta$, and the third panels show the obtained exponential cutoff $c^*$.

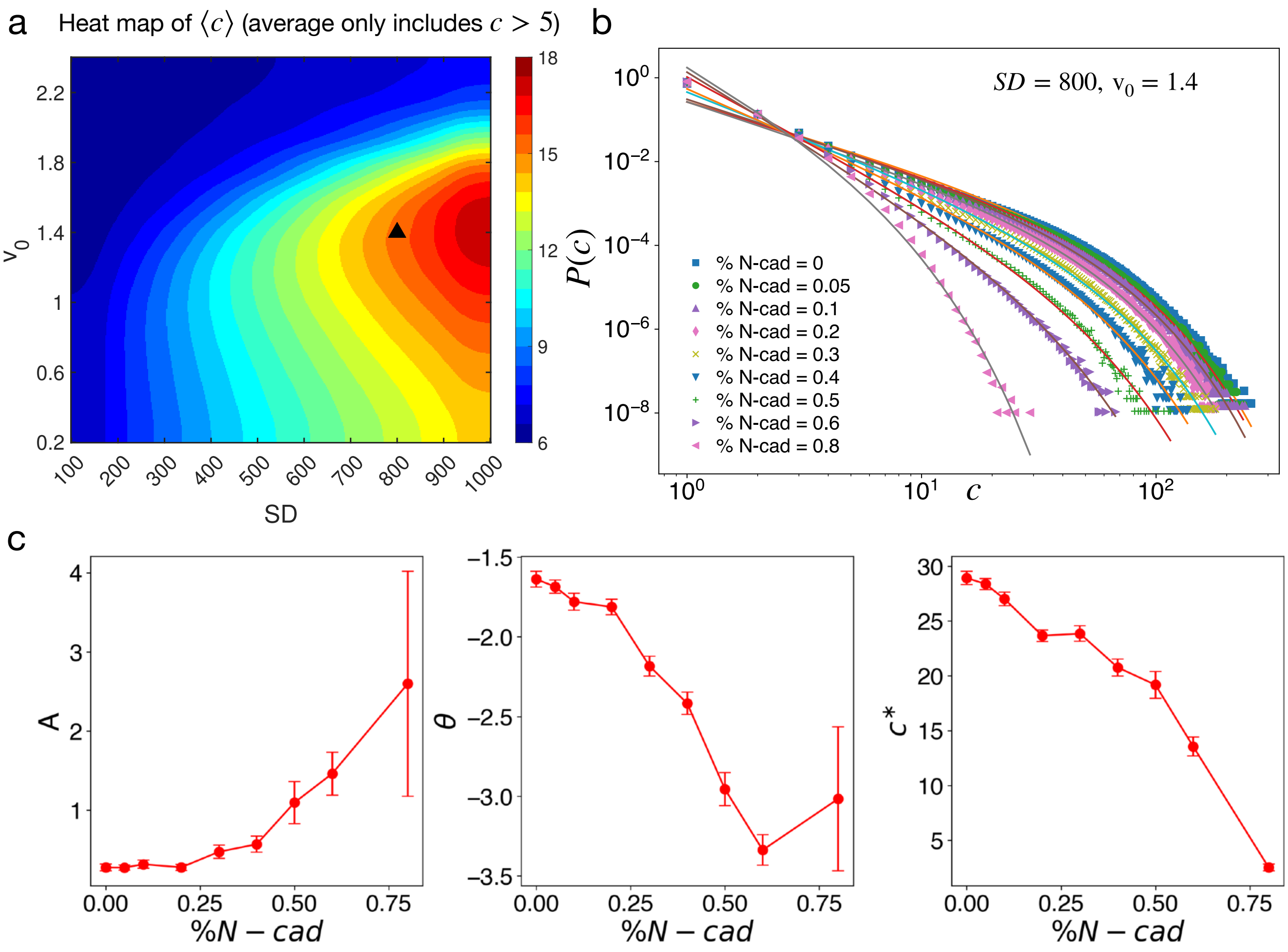


FIG. 3. (a) Average cluster size $\langle c \rangle$, computed excluding from monomers to the pentamers, of 100% E-cadherin junctions shown as a heatmap as a function of $v_0$ and $SD$. The triangle marks the conditions we choose to subsequently perform the cadherin isoform switch and the marked cluster size value is $c_{opt}$, the optimal threshold cluster size. (b) Evolution of $P(c)$ at increasing values of % N-cadherin at the junction. (c) Panels showing how the fitting parameters of $P(c)$ depend on % N-cadherin: from left, first panel shows the values of $A$, second panel shows $\theta$, and the third panel shows $c^*$.
